# Repurposing Pyramax® for the Treatment of Ebola Virus Disease: Additivity of the Lysosomotropic Pyronaridine and Non-Lysosomotropic Artesunate

**DOI:** 10.1101/2020.04.25.061333

**Authors:** Thomas R. Lane, Julie Dyall, Luke Mercer, Caleb Goodin, Daniel H. Foil, Huanying Zhou, Elena Postnikova, Janie Y. Liang, Michael R. Holbrook, Peter B. Madrid, Sean Ekins

## Abstract

We have recently identified three molecules (tilorone, quinacrine and pyronaridine tetraphosphate) which all demonstrated efficacy in the mouse model of infection with mouse-adapted Ebola virus (EBOV) model of disease and had similar *in vitro* inhibition of an Ebola pseudovirus (VSV-EBOV-GP), suggesting they interfere with viral entry. Using a machine learning model to predict lysosomotropism these compounds were evaluated for their ability to inhibit via a lysosomotropic mechanism *in vitro*. We now demonstrate *in vitro* that pyronaridine tetraphosphate is an inhibitor of Lysotracker accumulation in lysosomes (IC_50_ = 0.56 μM). Further, we evaluated synergy between pyronaridine and artesunate (Pyramax®), which are used in combination to treat malaria. Artesunate was not found to have lysosomotropic activity *in vitro* and the combination effect on EBOV inhibition was shown to be additive. Pyramax® may represent a unique example of the repurposing of a combination product for another disease.

## Introduction

The outbreaks of Ebola virus (EBOV) disease (EVD) in Africa have come at great human and financial cost (*1, 2*). For example, the outbreak in 2014-2016 killed over 11,000 and it is estimated that it resulted in $53bn in economic damage (*3*). The most recent outbreak in the Democratic Republic of the Congo, has killed more than 2200 people (*4*). Even with approval of a vaccine for prevention of EVD (*5*) there is still an urgent need to advance development of filovirus-specific antiviral therapeutics. A clinical trial (NCT03719586) investigated ZMapp (a monoclonal antibody cocktail) (*6*)), remdesivir (a small molecule), MAb114 (a monoclonal combination) (*7*)) and REGN-EB3 (monoclonal antibody combination) (*8*). These results showed that the antibodies REGN-EB3 and mAb114 had overall statistically similar survival rates of 71% and 66%, respectively. Unfortunately, ZMapp and remdesivir were less effective with a 51% and 47% survival rates, respectively (*9*). With remdesivir showing no statistically significant clinical efficacy the search for antiviral small molecules continues.

In an effort to repurpose drugs for the treatment of EVD, we have developed a Bayesian machine learning (ML) approach with a set of 868 anti-EBOV active molecules identified in a viral pseudotype entry assay and confirmed in an EBOV replication assay (*10, 11*). The EBOV ML model enabled us to virtually screen several thousand compounds and identify three active compounds against EBOV: tilorone, quinacrine and pyronaridine tetraphosphate (*12*). The three molecules inhibited EBOV in HeLa cells and demonstrated significant *in vivo* activity in the mouse-adapted EBOV (ma-EBOV) efficacy model (*13–16*) and in the guinea pig model of EBOV infection. The compounds also inhibited replication of multiple strains of EBOV and Marburg virus (MARV) (*17*). The trend for compounds to be active against both EBOV and MARV has been demonstrated before, with analysis of previously published data revealing an *in vitro* inhibition (IC_50_’s) correlation ((*10, 18*), Figure S1).

To date, we have not determined the mechanism of the antiviral compounds we have identified. Previously we evaluated pyronaridine, tilorone and quinacrine *in vitro* for its anti-EBOV activity (Zaire strain) in the type I IFN-deficient Vero 76 cell line (*19, 20*) and no antiviral activity was observed at any concentration below the 50% cytotoxicity concentration. In HeLa cells all three drugs demonstrated selectivity (*12, 14*). These observations support the hypothesis that their antiviral activity could be partially acting through or on the type I IFN-related innate immunity pathway (*15*). We also tested a combination of pyronaridine with tilorone in HeLa cells and evaluated the data with the BRAID model which suggested they are likely synergistic (*21*). Based on published data for tilorone and quinacrine, which are well known to be lysosomotropic agents, it was suspected that this may also be important. In addition, pyronaridine is used as an antimalarial in combination with artesunate (Pyramax®). We had previously determined that artesunate also has micromolar *in vitro* inhibitory activity against EBOV (*22*). We now assess whether pyronaridine accumulates in lysosomes and if there is any effect with artesunate or its active metabolite dihydroartemisinin against EBOV when combined with pyronaridine *in vitro*.

## Materials and Methods

### Chemicals and reagents

Pyronaridine tetraphosphate [4-[(7-Chloro-2-methoxybenzo[b][1,5]naphthyridin-10-yl)amino]-2,6-bis(1-pyrrolidinylmethyl)phenol phosphate (1:4)] (*12*) was purchased from BOC Sciences (Shirley NY). Tilorone was purchased from BOC Sciences. Quinacrine and Chloroquine were purchased from Cayman Chemicals (Ann Arbor, Michigan) and Sigma-Aldrich (St. Louis, MO), respectively. Artesunate was purchased from TRC Canada (North York, ON, Canada) and dihydroartemisinin (DHA) was purchased from Sigma-Aldrich (#D7439).

### NIAID screening

Pyronaridine tetraphosphate, tilorone and quinacrine were also tested (using the NIAID DMID services) against representatives of several viruses using human cells. The general methods have been described previously (*16*).

### Lysosomotropic machine learning model

The Assay Central software has been previously described (*23–32*) which uses the source code management system Git to gather and store structure-activity datasets collated in Molecular Notebook (Molecular Materials Informatics, Inc. in Montreal, Canada). The output is a high-quality dataset and a Bayesian model using extended-connectivity fingerprints of maximum diameter 6 (ECFP6) descriptors. Each model includes several metrics to evaluate and compare predictive performance as previously described in a relevant publication (*29*), including Receiver Operator Characteristic, Recall, Precision, F1 Score, Cohen’s Kappa (*33, 34*), and Matthews Correlation Coefficient (*35*). Applicability is representative of the overlap between the training and the test set. It is the quotient of the total number of ECFP6 fingerprints of the test molecule represented in the model divided by the total number of ECFP6 fingerprints of that test molecule. Generation and interpretation of prediction scores has been previously described (*36, 37*). The model consisted of curated data from a key paper from Nadanaciva *et al*. (*38*), where their quantitative approach to measuring lysosomotropic properties allowed for a direct activity threshold cut-off and was defined as an IC_50_ (decrease in LysoTracker Red staining) of ≥ 70 μM. A negative series of drugs that lack lysosomotropic properties from Kazmi *et al.* was also curated and added as inactive compounds (*39*) to the model.

### Lysosomotropic method

A previous published lysosomotropic assay by Nadanaciva *et al.* was used as the basis for the following work (*38*).

#### MCF7 cell culture conditions

The human metastatic mammary gland cell line MCF7 was obtained from American Type Culture Collection (ATCC# HTB-22). Cells were grown in Eagle’s minimum essential medium (Corning) supplemented with 10% fetal bovine serum (Gibco), 100 unit/ml penicillin and 100 μg/ml streptomycin (Corning) in a humidified incubator at 37°C and 5% CO_2_.

#### Lysosomotropic Assay

MCF7 cells were seeded into black walled clear bottom 96-well plates at 15,000 cells/well in 100 ul growth media and incubated for 48 hours (h). Cells were treated with drugs at 2-fold dilutions, with an initial testing concentration of 50 μM and an additional series of 9 tested dilutions (final 0.098 μM). Based on solubility restrictions, compounds for stocks were either solubilized in DMSO (tilorone, quinacrine, artesunate) or water (pyronaridine, chloroquine). Control wells included cells treated with DMSO or water. To start assay, 0.5 μl of appropriate compound stock or control was added using Biomek NX^p^ (Beckman Coulter) and incubated at 37°C, 5% CO_2_ for 3 hours. LysoTracker Red (75 nM) (ThermoFisher) was then added and incubated for 30 min followed by a wash in phosphate buffered saline (PBS). The cells were immediately fixed with 10% formalin at room temperature for 15 min. Cells were then stained with Hoechst (5 μg/ml) (Sigma-Aldrich) in PBS for 10 min at room temperature. Following cell staining they were washed in PBS. Each experimental run tested a series of compounds in triplicate and was repeated on two different days (n=6 for each compound series) with multiple DMSO (n=12) and water (n=24) controls per plate.

Imaging was done using a CellInsight CX5 High Content Screening Platform (Thermo Scientific) with 10X objective. Fluorescence was measured with Hoechst (nuclei) and LysoTracker Red (lysosomes) in channel 1 and 2, respectively. A total of 3 to 4 fields were captured for all wells. For analysis, nuclei were identified, and a circular mask was extended out 5 pixels to represent the cell. Total intensity of the fluorescent signal from Lysotracker Red within the mask area was then used to represent the lysosomal staining in the cells. Data was normalized to controls and then analyzed with GraphPad Prism version 8.00. Error bars of dose-response curves represent the SEM of the replicates.

### Cell Viability

MCF7 cells were seeded in white walled clear bottom 96-well plates at 15,000 cells/well in 100 μl growth media and incubated for 48 h. To start the assay, 0.5 μl of compound stock or control was added using Biomek NX^p^ and incubated at 37°C, 5% CO_2_ for 3.5 hours. Following compound incubation, 80 μl of CellTiter-Glo (Promega) was added. The plates were shaken on an orbital shaker at 300 rpm for 20 min and then read on an Envision 2104 Multilabel reader (PerkinElmer). Experiments were repeated in triplicate and data was analyzed with Graphpad Prism version 8.00. Error bars of dose-response curves represent the SEM of the replicates.

### *In vitro* combination studies methods

The *in vitro* infection inhibition of EBOV/Mak (Makona, IRF0165, 1.98E7 PFU/mL) was performed in HeLa and Huh 7 cells. HeLa cells were seeded at 3 × 10^4^ cells/well in 96-well plates. After 24 hr the drugs were added to cells in a 6×6 matrix with 2-fold serial dilutions with a starting concentration of 30 μM. The experiment was run on 3-4 replicate plates. The experiment was run on 2 different days. Cells were infected with virus 1 h after the addition of the drugs in BSL4-containment at a multiplicity of infection (MOI) of 0.21 (Huh 7) or 0.5 (HeLa). After 48 h, plates were fixed and virus was detected with a mouse antibody specific for EBOV VP40 protein (#B-MD04-BD07-AE11, made by US Army Medical Research Institute of Infectious Diseases, Frederick MD under Centers for Disease Control and Prevention contract) (*40*) followed by staining with anti-mouse IgG-peroxidase labeled antibody (KPL, Gaithersburg, MD, #074-1802). Luminescence was read on an Spark 20M plate reader (Tecan US, Morrisville, NC). The signal of treated, infected wells was normalized to uninfected control wells and measured (in percent) relative to untreated infected wells. Non-linear regression analysis was performed, and the 50% inhibitory concentrations (EC_50_s) were calculated from fitted curves (log [agonist] versus response [variable slope] with constraints to remain above 0% and not exceed 100%) (GraphPad Software version 8.0, La Jolla, CA). The EBOV drug screen assay was performed with three replicates for each drug concentration. Error bars of dose-response curves represent the SEM of the replicates. For quantitation of drug toxicity, HeLa cells were mock infected (no virus) and treated with drug dilutions under the same conditions as the infected cells. After 48 h, cell viability was measured using the CellTiter Glo Luminescent Cell Viability Assay kit according to manufacturer’s protocol (Promega, Madison, WI).

### Combination Analysis using BRAID and SynergyFinder

The BRAID analysis (*21*) service calculates synergy by fitting data to a seven-variable function. The variable κ represents a quantitative synergy value where κ□<□0 implies antagonism, κ□=□0 implies additivity, and κ□>□0 implies synergy. As an additional reference, “strong synergy” corresponds to κ□=□2.5, “mild synergy” corresponds to κ□=□1, “mild antagonism” corresponds to κ□=□−0.66, and “strong antagonism” corresponds to κ□=□−1. To assess if the combined inhibitory effect of pyronaridine and or artesunate/dihydroartemisinin on EBOV was synergistic, additive, or antagonistic we analyzed a 6×6 checkboard assay with these pairs of drugs at various combined concentrations in HeLa and Huh 7 cells. It is noted that inhibition data under toxic concentrations (consistently >50% cell death) were removed from the analysis. Inclusively, this consisted of only individual and combined experiments with concentrations of pyronaridine that exceeded its CC_50_ (i.e. 5 μM concentrations in HeLa cells only). All toxicity data was retained for BRAID analysis.

The SynergyFinder analysis service (*41*) similarly calculates the degree of combination, synergy or antagonism by comparing the observed drug combination response against the expected response, while assuming there is no interaction between the two drugs. These scores were calculated using the Loewe reference additivity mode (*42*). The threshold to define a good synergy score is variable, but the program developers suggest that synergy scores near 0 gives limited confidence on synergy or antagonism and a score < −10 or >10 are expected to be antagonist or synergistic, respectively.

## Results

### NIAID invitro screening

Pyronaridine, quinacrine and tilorone were previously demonstrated to be active against EBOV in HeLa cell but not Vero cells (*14*). We have now tested these compounds against Adenovirus 5, Human papillomavirus 11, Chikungunya virus, Dengue virus 2, Powassan virus, Rift valley virus, Yellow Fever virus and human cytomegalovirus in additional human cells through the use of NIAID screening resources. None however showed selectivity in the cell lines and concentrations tested, but this may be due to high cytotoxicity and or the insufficient range of concentrations tested in these cell lines (Table S1A-C)

### Lysosomotropic machine learning model predictions

A Bayesian machine learning model with 52 compounds (23 were classed as lysosomotropic) was generated from published data using Assay Central™ with 5-fold cross validation ROC = 0.765 (Figure 1). Additional model statistics suggest that the model is potentially useful for scoring compounds to predict lysosomal accumulation (Table 1). Tilorone, pyronaridine and artesunate were used as a prospective test set. All compounds were correctly predicted, with tilorone and pyronaridine predicted to be lysosomotropic, while artesunate was predicted to not be lysosomotropic.

**Table 1.** Physicochemical properties and Assay Central lysosomotropic machine learning predictions for compounds tested *in vitro*. An applicability score of 1 indicates that all the fragments are in the model and may indicate the molecule is in the training set (chloroquine is in the training set). (Calculated with ACD/Labs PhysChem Batch program^$^, (*59*)) Predicted pka’s (negative log of the acid dissociation constant) were obtained from drugbank, which were initially calculated using Chemaxon. AlogP (predicted log octanol-water partition coefficient was calculated via Discovery Studio).

| **Name** | **pK<sub>a</sub> (predicted)** | **Pka (Experimental)** | **AlogP** | **Lysosomotropic Prediction Score** | **Lysosomotropic Applicability Score** |
| --- | --- | --- | --- | --- | --- |
| Chloroquine | 10.32 (Strongest Base) | 4.0, 8.4 and 10.2 (60) | 4.34 | 1.09 | 1 |
| Artesunate | 3.77 (Strongest Acid), -4.2 (Strongest Base) | 4.6 (61) | 1.84 | 0.31 | 0.21 |
| Quinacrine | 10.33 (Strongest Base) | N/A | 5.67 | 1.00 | 0.68 |
| Tilorone | ~8.6<sup>\$</sup> | N/A | 4.56 | 0.75 | 0.69 |
| Pyronaridine | 7.96 (Strongest Acid), 10.08 (Strongest Base) | 7.08, 7.39, 9.88 and 10.30 (62) | 6.19 | 0.68 | 0.51 |

**Figure 1.**
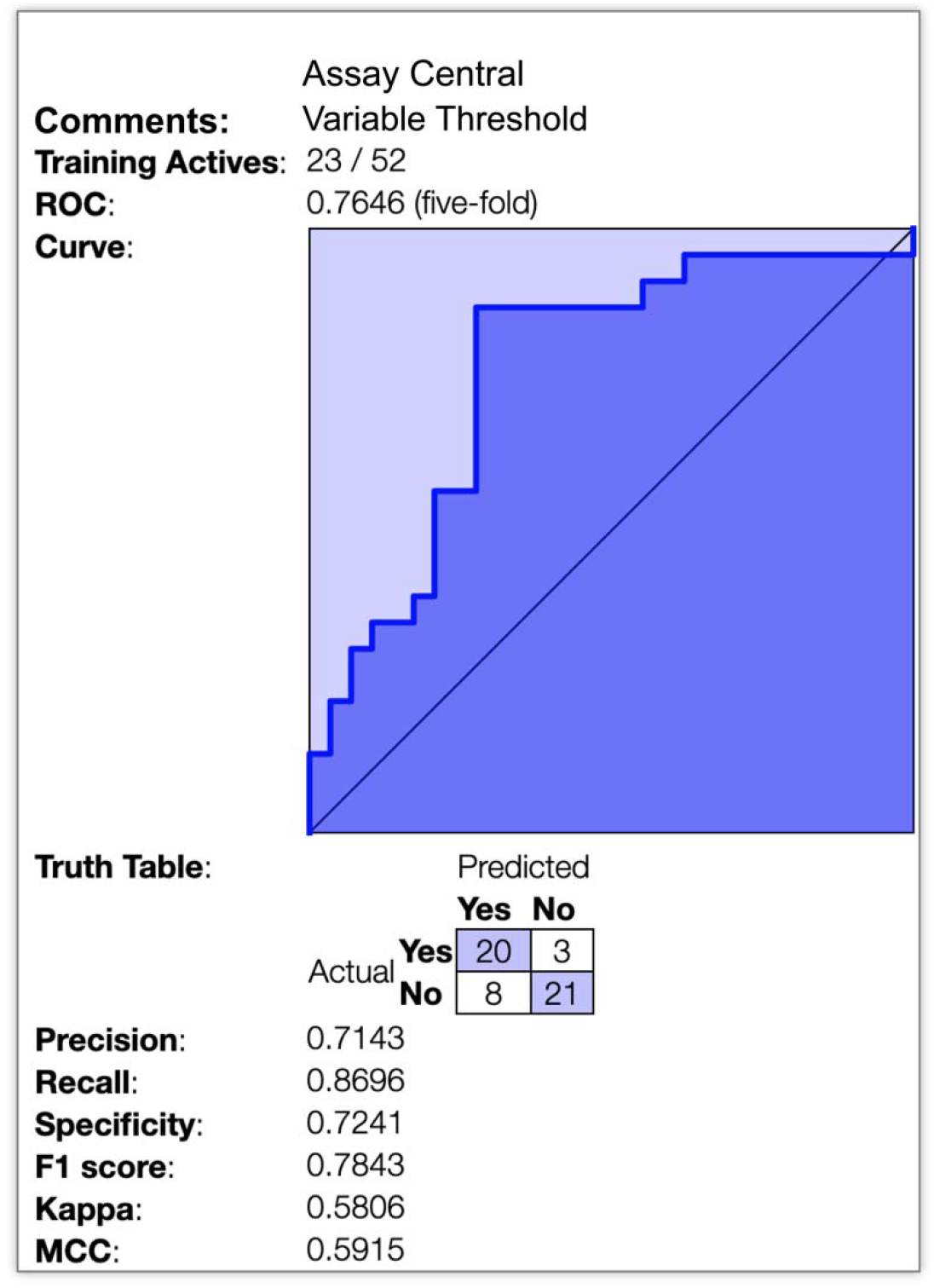
Lysosomotropic machine learning model. 5-fold cross validation receiver operator curve as well as multiple metrics depicting the internal validation of this Bayesian model (ECFP6).

### *In vitro* inhibition of lysosomal accumulation of Lysotracker

Pyronaridine tetraphosphate was found to be a potent inhibitor of Lysotracker accumulation in MCF7 lysosomes *in vitro* (IC_50_ = 0.56 μM). In contrast, artesunate showed no appreciable inhibition of Lysotracker (Figure 2). Tilorone (IC_50_ = 3.09 μM), chloroquine (IC_50_ = 6.21 μM) and finally quinacrine (IC_50_ = 7.31 μM) were less potent inhibitors of Lysotracker accumulation in MCF7 lysosomes *in vitro* (Figure S2).

**Figure 2.**
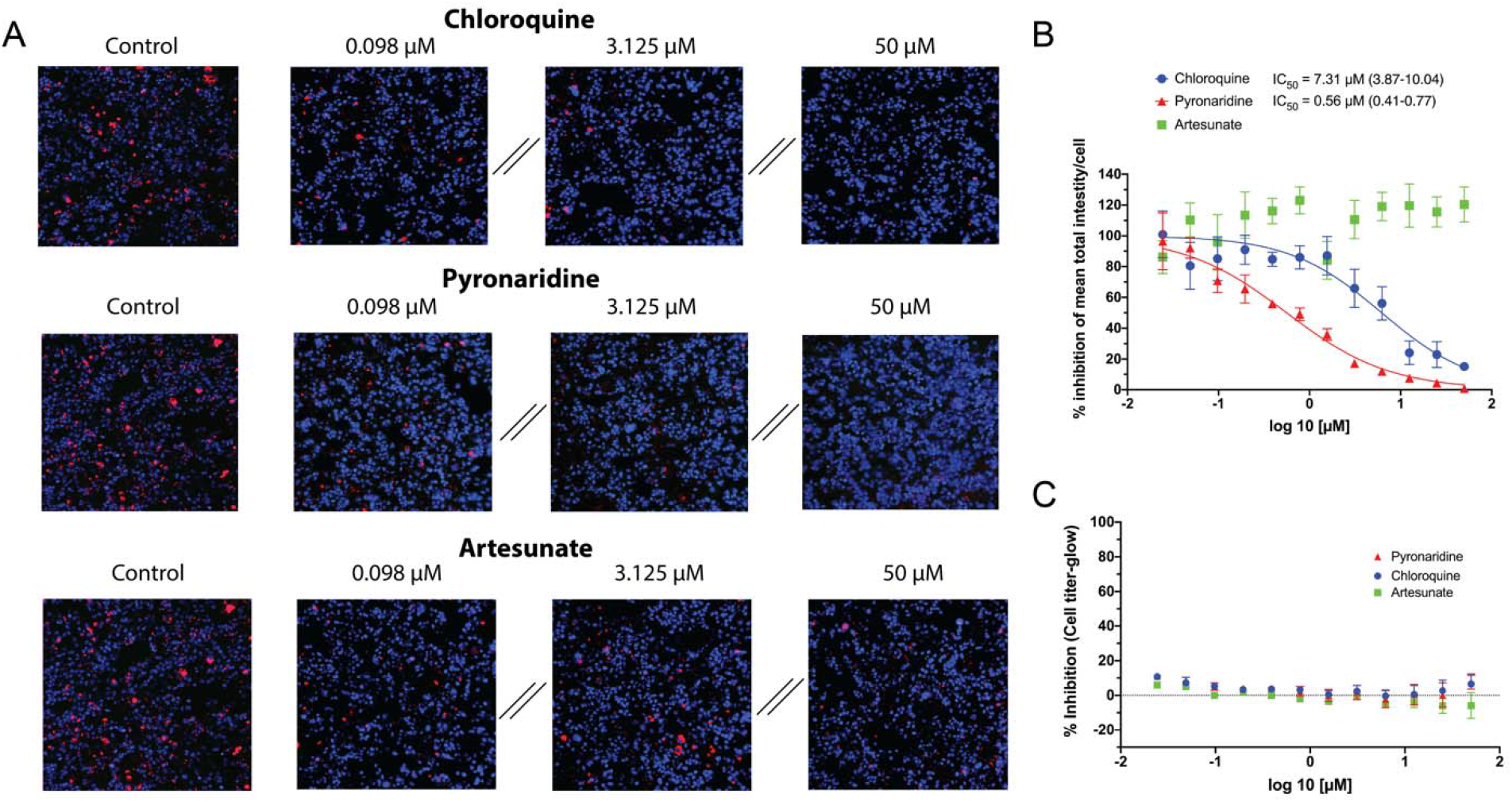
Inhibition analysis of total fluorescent intensity/cell of lysotracker red by chloroquine, pyronaridine and artesunate in MCF7 Cells. Lysotracker accumulation in lysosomes is pH dependent, therefore a reduction in signal from the lysotracker suggests a pH increase in these organelles. This is proposed to be caused by accumulation of the charged base of the lysosomotropic compound in the lysosome, which in a lower pH environment becomes neutralized and trapped in the organelle. A) Representative images showing Lysotracker lysosomal accumulation inhibition at various concentrations. B) Graphical representation and quantification (Parentheses represent 95% CI) of the dose-dependent effect of on Lysotracker accumulation in lysosomes (Error bars represent SEM). Outliers were identified using the ROUT method (Q=10%) and consequentially removed. C) Measure of cellular toxicity at concentrations and times mimicking the inhibition assays.

### Combination Analysis

The BRAID analysis (Figure 3) of pyronaridine and artesunate *in vitro* inhibition data from the checkerboard assay indicates additivity of these molecules in HeLa cells. Artesunate ameliorates the toxicity of pyronaridine in the checkboard assay and therefore indirectly potentiates pyronaridine. The non-linear regression, 4-parameter curve fit (Hill equation) for the artesunate control in Huh 7 cells suggested a plateau at ~60% inhibition (Figure S3), therefore the combination data for this cell-line and pyronaridine/artesunate pair was not included in the analysis. Pyronaridine and DHA similarly shows an additive effect in HeLa and Huh 7 cells, both with a parallel reduction in toxicity based on the BRAID analysis (Figure S4). A secondary analysis using Synergyfinder (Figures S5-S6) also suggests that this combination indirectly potentiates pyronaridine in HeLa cells, but in Huh 7 cells these interpretations are ambiguous (Figure S7).

**Figure 3.**
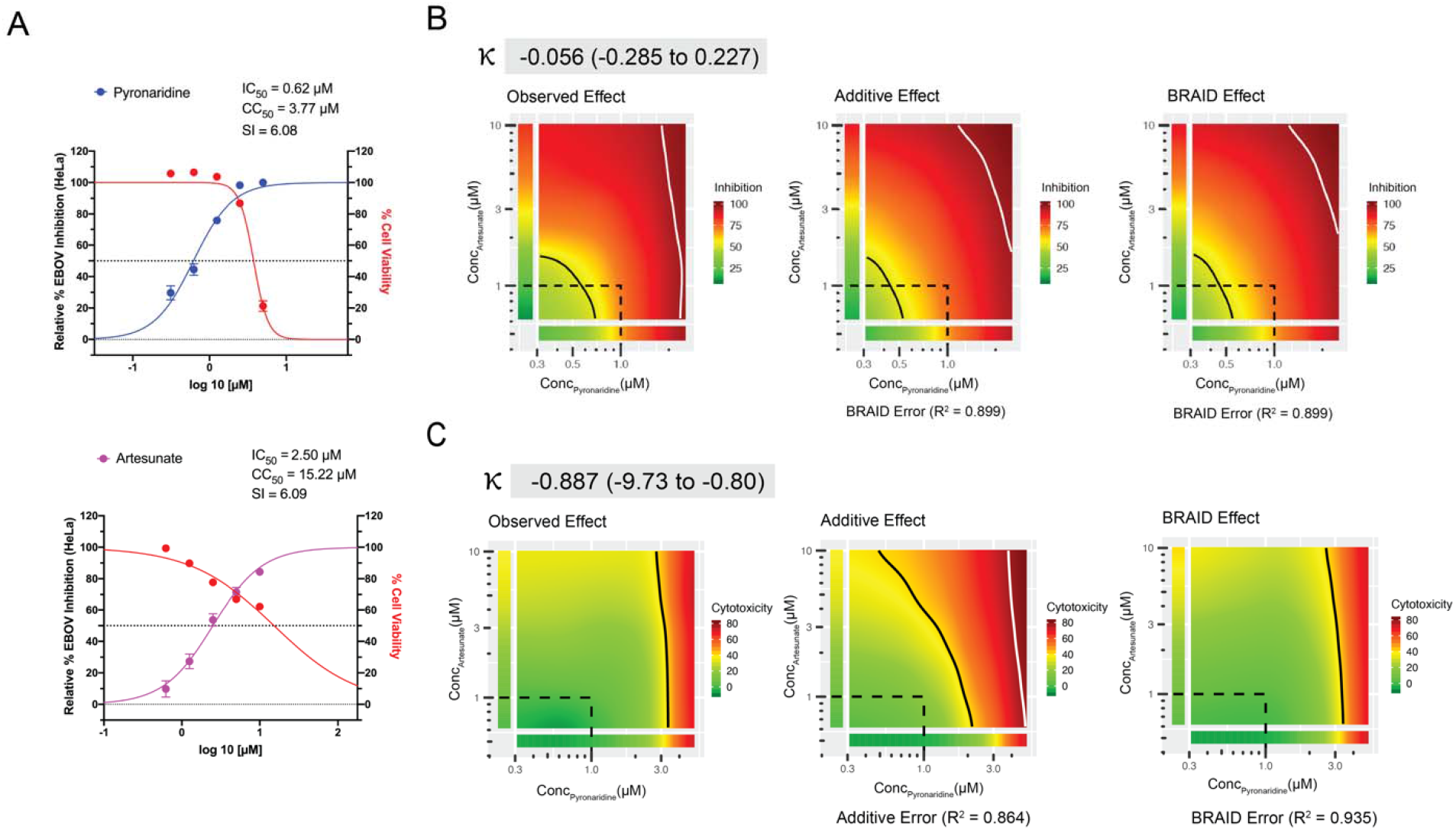
Combination data for the pyronaridine and artesunate checkerboard assay in HeLa cells. A) Inhibition/cytotoxicity plots for the pyronaridine and artesunate controls (compound tested in the absence of the other compound). Controls were run in triplicate at 5 concentrations per plate, so the total number from replicates for each compound varied (Pyronaridine, n=27; Artesunate, n=18). Error bars represent the SEM at each concentration tested. B) Graphical representations (from left to right) of the inhibition plots of the smoothed raw data, predicted additive inhibition and predicted inhibition using the 7-parameter BRAID analysis. It is noted that inhibition data under toxic concentrations (>50% cell death) were removed from the analysis. The “Additive” or “BRAID” error represents the corresponding accuracy of fit with the “Observed Effect”. κ represents the combinatory effect where “strong synergy” corresponds to κ□=□2.5, “mild synergy” corresponds to κ□=□1, “mild antagonism” corresponds to κ□=□−0.66, and “strong antagonism” corresponds to κ□=□−1. C) Representation of the cytotoxicity (toxicity is representative of % cell death from control) arranged in the same manner as inhibition.

## Discussion

Within the last 5 years we have seen two major EVD outbreaks in Africa. These led to renewed efforts to develop treatments for this virus. The actives and inactives of several *in vitro* high throughput drug screens (*10, 18, 43*) have been used to develop computational models for predicting anti-EBOV activity of compounds. More recently, combinations of approved drugs found in these and other studies have suggested synergistic combinations (*44–47*). To date, none of these many efforts for EBOV have resulted in a clinical antiviral candidate. Several small molecule antivirals were felled at the hurdle of animal models, specifically the transition from mouse to the guinea pig model. Compounds that have failed to show *in vivo* efficacy against EBOV following this well-trodden route include chloroquine (*18, 48*), azithromycin (*18*), amiodarone (*49*), BGB324 (*50*), NCK8 (*50*) and 17-DMAG (*50*). As discussed previously (*17*), this may be due to differences in drug metabolism making the model inappropriate for EBOV.

Efforts to improve the efficiency and cost effectiveness of EBOV drug discovery have involved our efforts to identify several molecules using ML which have progressed through *in vitro* and *in vivo* testing (*12–15, 17, 51*). We have also previously used these ML models to predict *in vitro* efficacy for drugs that were then tested against EBOV (*12, 22*). The mechanism for these three compounds against EBOV is unknown. Others have demonstrated that compounds with physicochemical properties such as a basic pKa (> 6.5) and cLogP of > 2 tend to be lysosomotropic (*38*) and they accumulate in the lysosomes. We have now taken an ML approach to predict potential for a lysosomotropic mechanism using published *in vitro* data (*38, 39*) along with ECFP6 molecular fingerprints and a Bayesian algorithm. We have now performed several *in vitro* studies to validate the predictive ability of this model as well as infer the potential mechanism of one of these EBOV drugs. Pyronaridine clearly is a potent lysosomotropic agent, more so than all the other molecules tested. There is a strong correlation between published anti-EBOV activity and the lysosomotropic property (Table S2) for a large number of drugs. All of the compounds that were considered actives in our model were researched to identify whether they had been previously tested against EBOV and or MARV either with a psuedovirus/VLP and or a competent virus inhibition assay. 21 of 23 of these compounds had been tested previously and all inhibited these viruses, with AC_50_s almost all in the nM to low μM range. This is certainly not a comprehensive list of all lysosomotropic compounds, but this strongly supports the notion that the lysosomotropic characteristic is directly related to the antiviral activity of the compounds within this model. Artesunate which is also similarly active against EBOV was found not to share this physiological characteristic. The initial combination of these two drugs to form Pyramax® was to avoid drug resistance of *Plasmodium* parasites, the causative agents of Malaria and has been extensively reviewed (*52*). The combinations of these two molecules are additive in inhibiting EBOV replication *in vitro* but with a reduced cytotoxicity as compared to the individual treatments (Figure 2 and 3). Previous work has suggested that it is possible to identify pairs of drugs that block EBOV infection *in vitro* via the same methodology as used here (*47*) and these prior data have been used with other software to suggest a variation in prioritizing drug pairs based on selective efficacy (*53*), which considers both synergy and toxicity. This software independently confirms our observations with the BRAID analysis (Figure S5-7).

This current study has implications outside of EBOV, with the increased interest in antivirals for testing against SARS-CoV-2, and in particular the heavy focus on chloroquine and hydroxychloroquine, which have shown low micromolar activity *in vitro* against this virus (*54–57*). Pyronaridine has recently also been shown to have some limited activity against SAR-CoV-2 in Vero cells (*54*), but Vero cells may not be as appropriate as human cells to test compounds such as pyronaridine (low *in vitro* selectivity index, but high *in vivo* antiviral activity). Based on our previous findings, this leaves the distinct possibility that there has been an underestimate of pyronaridine’s antiviral inhibition potential. Our current data also suggests that we should be testing artesunate versus additional viruses as well as in combination with pyronaridine.

Frequently, single drugs are repurposed for new uses (*58*), to our knowledge there is no precedent for a two drug combination being repurposed for the same indication. We have previously estimated that the dose used for treating malaria patients may have a beneficial effect in EBOV patients (*14*), further indicating the potential for direct repurposing without the need to change dose, route or formulation. From our experience and insights gained during the discovery of the antiviral properties of pyronaridine, tilorone and quinacrine we propose our ML approach could be optimized by adding the additional ML model for the lysosomotropic mechanism described here. This would enable us to create a computational pipeline to identify new antivirals more rapidly that could have this lysosomotropic property and hence direct the antiviral mechanism of action investigations. Molecules with this mechanism may also have more utility as broad-spectrum antivirals which is needed to counter flare ups of viruses like EBOV, MARV or potential pandemics such as SARS-CoV-2.

## ACKNOWLEDGMENTS

Dr. Mindy Davis is gratefully acknowledged for assistance with the NIAID virus screening capabilities, Task Order number B22. We acknowledge numerous discussions with Dr. Jason E. Comer, Dr. Shane Massey, Dr. Manu Anantpadma, Dr. Robert A. Davey and Dr. Joel S. Freundlich.

