## Supplemental files for "Repurposing Pyramax® for the Treatment of Ebola Virus Disease: Additivity of the Lysosomotropic Pyronaridine and Non-Lysosomotropic Artesunate"

^c^Cambrex, 3501 Tricenter Blvd, Suite C ∙ Durham, NC 27713, USA.

^d^SRI International, 333 Ravenswood Avenue, Menlo Park, CA 94025, USA.

**Short running title:** Pyronaridine combination treatment for Ebola

**Keywords:** Antiviral, Ebola, Lysosomotropic, Machine learning

Table S1. NIAID *in vitro* virus testing of a. pyronaridine, b. quinacrine and c. tilorone in human cells.

a.

|  | **Cell line** | **EC_50_ (µM)** | **EC_90_ (µM)** | **CC_50_ (µM)** | **SI_50_** | **SI_90_** |
| --- | --- | --- | --- | --- | --- | --- |
| Adenovirus 5 | HFF | >6 | >6 | 15.75 | <3 | <3 |
| Human papillomavirus 11 | C-33A | >6 | >6 | 16.48 | <3 | <3 |
| Chikungunya virus | Huh7 | >2.4 |  | 2.4 | 0 |  |
| Dengue virus 2 | Huh7 | >3.2 |  | 3.2 | 0 |  |
| Powassan virus | BHK-21 | >3.2 |  | 3.2 | 0 |  |
| Rift Valley virus | Huh7 | >2.4 |  | 2.4 | 0 |  |
| Yellow Fever virus | Huh7 | >3.2 |  | 3.2 | 0 |  |
| Human cytomegalovirus | HFF | >6 | >6 | 17.23 | <3 | <3 |

b.

|  | **Cell line** | **EC_50_ (µM)** | **EC_90_ (µM)** | **CC_50_ (µM)** | **SI_50_** | **SI_90_** |
| --- | --- | --- | --- | --- | --- | --- |
| Adenovirus 5 | HFF | >1.2 | >1.2 | 3.98 | <3 | <3 |
| Human papillomavirus 11 | C-33A | >6 | >6 | 15.46 | <3 | <3 |
| Chikungunya virus | Huh7 | >3.2 |  | 3.2 | 0 |  |
| Dengue virus 2 | Huh7 | >3.2 |  | 3.2 | 0 |  |
| Powassan virus | BHK-21 | >3.2 |  | 3.2 | 0 |  |
| Rift Valley virus | Huh7 | >1.9 |  | 1.9 | 0 |  |
| Yellow Fever virus | Huh7 | >3.2 |  | 3.4 | 0 |  |
| Human cytomegalovirus | HFF | >6 | >6 | 17.09 | <3 | <3 |

c.

|  | **Cell line** | **EC_50_ (µM)** | **EC_90_ (µM)** | **CC_50_ (µM)** | **SI_50_** | **SI_90_** |
| --- | --- | --- | --- | --- | --- | --- |
| Adenovirus 5 | HFF | >6 | >6 | 15.06 | <3 | <3 |
| Human papillomavirus 11 | C-33A | >6 | >6 | 20.71 | <3 | <3 |
| Chikungunya virus | Huh7 | >0.1 |  | <0.1 | 0 |  |
| Dengue virus 2 | Huh7 | >3.2 |  | 3.2 | 0 |  |
| Powassan virus | BHK-21 | >3.2 |  | 3.2 | 0 |  |
| Rift Valley virus | Huh7 | >10 |  | 10 | 0 |  |
| Yellow Fever virus | Huh7 | >3.2 |  | 3.2 | 0 |  |
| Human cytomegalovirus | HFF | >6 | >6 | 29.35 | <5 | <5 |

Table S2. Antiviral properties of the compounds designated as lysosomotropic (based on various criteria) against EBOV and or MARV.

| **Drug name** | **Decrease in LysoTracker_ Red staining, IC_50_**  **(µM)** | **EBOV** | **Pseudovirus**  **EBOV** | **MARV** | **Pseudovirus MARV** | **Block EBOV VLP entry** |
| --- | --- | --- | --- | --- | --- | --- |
| Amiodarone | 3.8 ± 2  (*38*) | 0.387 µM (1.68)  (*62*)  7.6  (*17*) | 0.81  (*10*) 2.02  (*62*) | NT | 2.36  (*10*)1.73  (*62*) | 4.43  (*63*) |
| Aripiprazole | 19.3 ± 5  (*38*) | 8.1 (0.42)- 3.76 (0.18)  (*64*) | NT | NT | NT | NT |
| Astemizole | 0.95 ± 2  (*38*) | 6.17 (1.34)- 1.37 (0.038)  (*64*) | NT | NT | NT | NT |
| Bepridil | 14.5 ± 4  (*38*) | 5.08 (0.38)- 3.21 (0.15)  (*64*) | 23.68  (*10*) | NT | 47.22  (*10*) | NT |
| Chloroquine | 11.1 ± 2  (*38*) | NT | 1.33  (*10*) | NT | 6.4  (*10*) | 15.3  (*63*) |
| Chlorpromazine | 5.8 ± 2  (*38*) | Only tested at single concentration, but significant at 30 µM  (*65*) | 1.96  (*10*) | NT | 7.45  (*10*) | NT |
| Clomipramine | 12.4 ± 2  (*38*) | 11.4 (0.15)- 2.57 (0.16)  (*64*) | NT | NT | NT | 4.99  (*63*) |
| Desipramine | 4.6 ± 2  (*38*) | 3-5 µM  (*66*) | NT | NT | NT | NT |
| Fluoxetine | 6.5 ± 2  (*38*) | NT | 1.71  (*10*) | NT | 4.21  (*10*) | NT |
| Imipramine | 10.7 ± 5  (*38*) | NT | 3.84  (*10*) | NT | 8.6  (*10*) | 13.7  (*63*) |
| Lapatinib | 17.3 ± 2  (*38*) | NT | NT | NT | NT | 0.398  (*63*) |
| Nortriptyline | 10.6 ± 5  (*38*) | NT | 10µM had 83.1% inhibition (16.6% toxicity)  (*10*) | NT | 10µM had 61.5% inhibition  (*10*) | NT |
| Paroxetine | 5.3 ± 2  (*38*) | 7.45 (0.41)- 1.38 (0.076)  (*64*) | NT | NT | NT | NT |
| Promethazine | 8.7 ± 2  (*38*) | 19.4  (*67*) | NT | 19.1  (*67*) | NT | NT |
| Sertraline | 6.6 ± 3  (*38*) | 3.13 (0.24)- 1.44 (0.057)  (*64*) | NT | NT | NT | 2.73  (*63*) |
| Sunitinib | 5.6 ± 2  (*38*) | NT | 0.47 (*45*) | NT | NT | 1.91  (*63*) |
| Tamoxifen | 3.8 ± 2  (*38*) | NT | NT | NT | NT | 0.734  (*63*) |
| Thioridazine | 1.5 ± 3  (*38*) | 6.24 (0.79)- 2.06 (0.12)  (*64*) | NT | NT | NT | NT |
| Gefitinib | NT | NT | NT | NT | NT | NT |
| Dasatinib | NT | 4.23 (0.20)-16.5 (4.6) (*64*) | NT | NT | NT | NT |
| Propranolol | NT | NT | NT | NT | NT | NT |
| Nicardipine | NT | NT | NT | NT | NT | NT |
| Imatinib | NT | 2.38±1.49 (*40*) | NT | NT | NT | NT |

Figure S1. Correlation of *in vitro* MARV and EBOV activity for compounds from previous *in vitro* screens (*10, 17*).

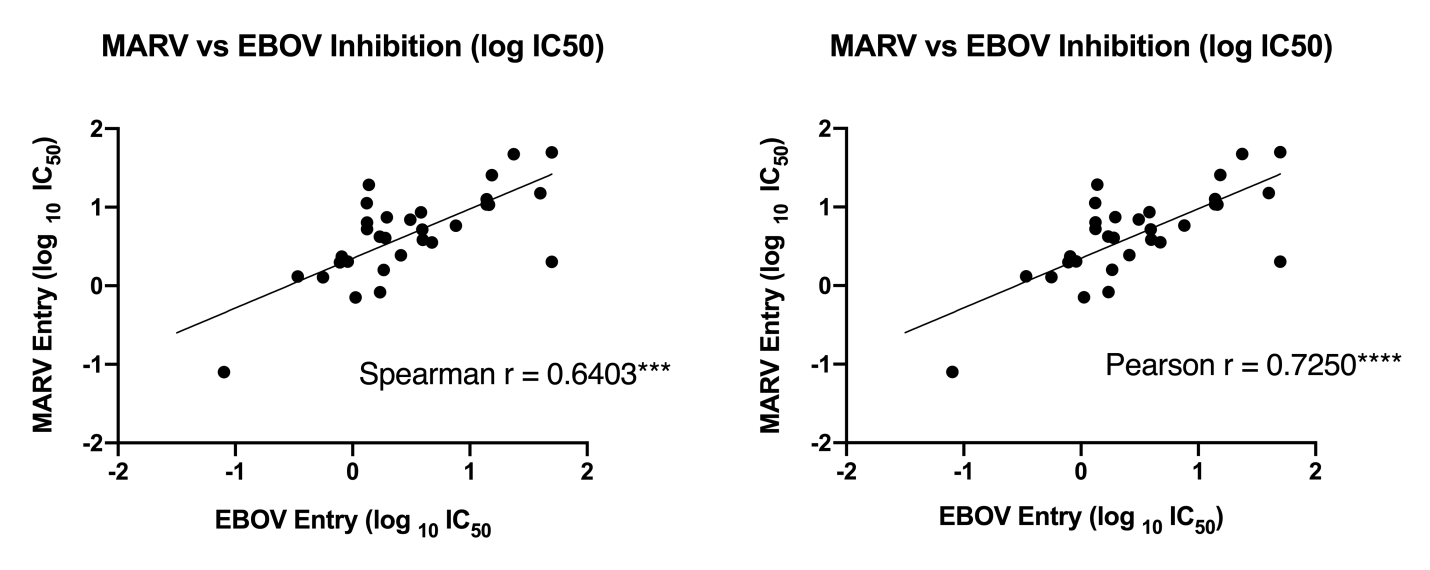

Figure S2. Inhibition of total fluorescent intensity/cell of lysotracker red in MCF7 Cells by chloroquine, quinacrine and tilorone. Lysotracker accumulation in lysosomes is pH dependent, therefore a reduction in signal from the lysotracker suggests a pH increase in these organelles. This is proposed to be caused by accumulation of the charged base of the lysosomotropic compound in the lysosome, which in a lower pH environment becomes neutralized and trapped in the organelle. A) Representative images showing Lysotracker lysosomal accumulation inhibition at various concentrations. B) Graphical representation and quantification (Parentheses represent 95% CI) of the dose-dependent effect of on Lysotracker accumulation in lysosomes (Error bars represent SEM). Outliers were identified using the ROUT method (Q=10%) and consequentially removed. C) Measure of cellular toxicity at concentrations and times mimicking the inhibition assays.

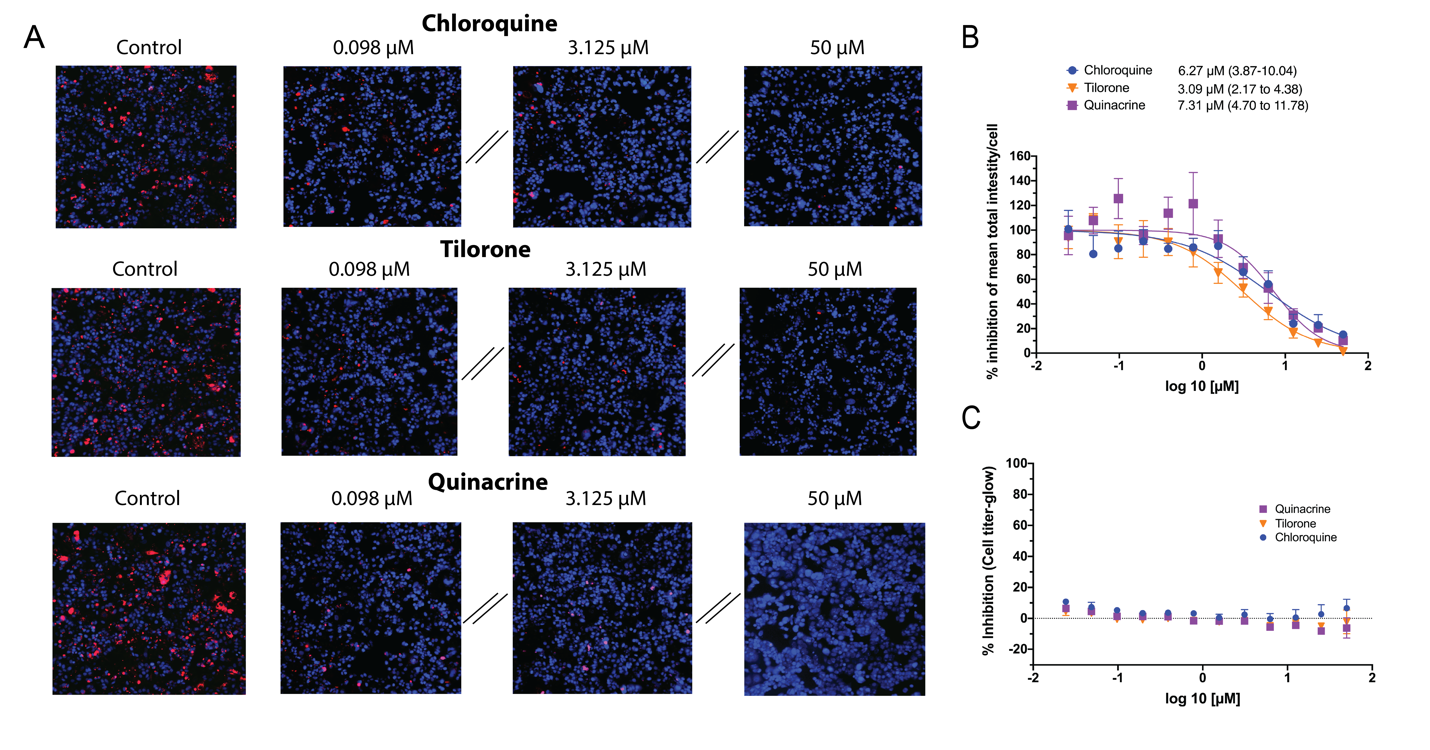

Figure S3. Fitted inhibition EBOV data for artesunate in Huh7 cells. A) Best fit with no max inhibition constraints (max inhibition plateau ~60%; IC50 = 1.548 (95% CI = 1.337-1.829)) and B) best fit with a forced max plateau of 100% inhibition (IC50 = 4.18 (95% CI = 3.634 – 4.874)). Due to the ambiguity of the max inhibition of artesunate against EBOV in Huh7 cells, these data were not included in the synergy analysis.

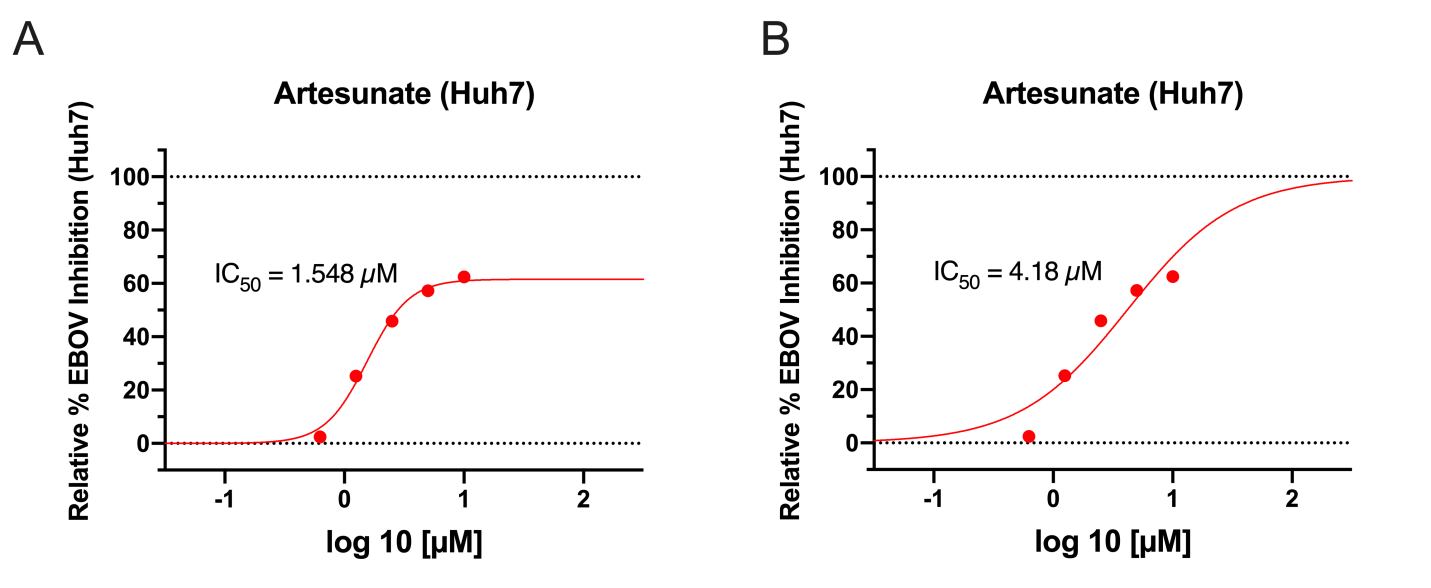

Figure S4. Combination data for the pyronaridine and DHA checkerboard assay in HeLa (A-C) and Huh7 (D-F) cells. A and F are the inhibition/cytotoxicity plots for the controls (compound tested in the absence of the other compound) in these two different cell lines. Controls were run in triplicate at 5 concentrations per plate, so the total number from replicates for each compound varied (Pyronaridine, n=27; DHA, n=9). Error bars represent the SEM at each concentration tested. B and E are graphical representations (from left to right) of the inhibition plots of the smoothed raw data, predicted additive inhibition and predicted inhibition using the 7-parameter BRAID analysis. It is noted that inhibition data under toxic concentrations (>60% cell death) were removed from the analysis. The Additive or BRAID error represents the corresponding accuracy of fit with the Observed Effect. κ represents the combinatory effect where “strong synergy” corresponds to κ = 2.5, “mild synergy” corresponds to κ = 1, “mild antagonism” corresponds to κ = −0.66, and “strong antagonism” corresponds to κ = −1. C and F mirror the inhibition organization but are representative of cytotoxicity (toxicity is representative of % cell death from control).

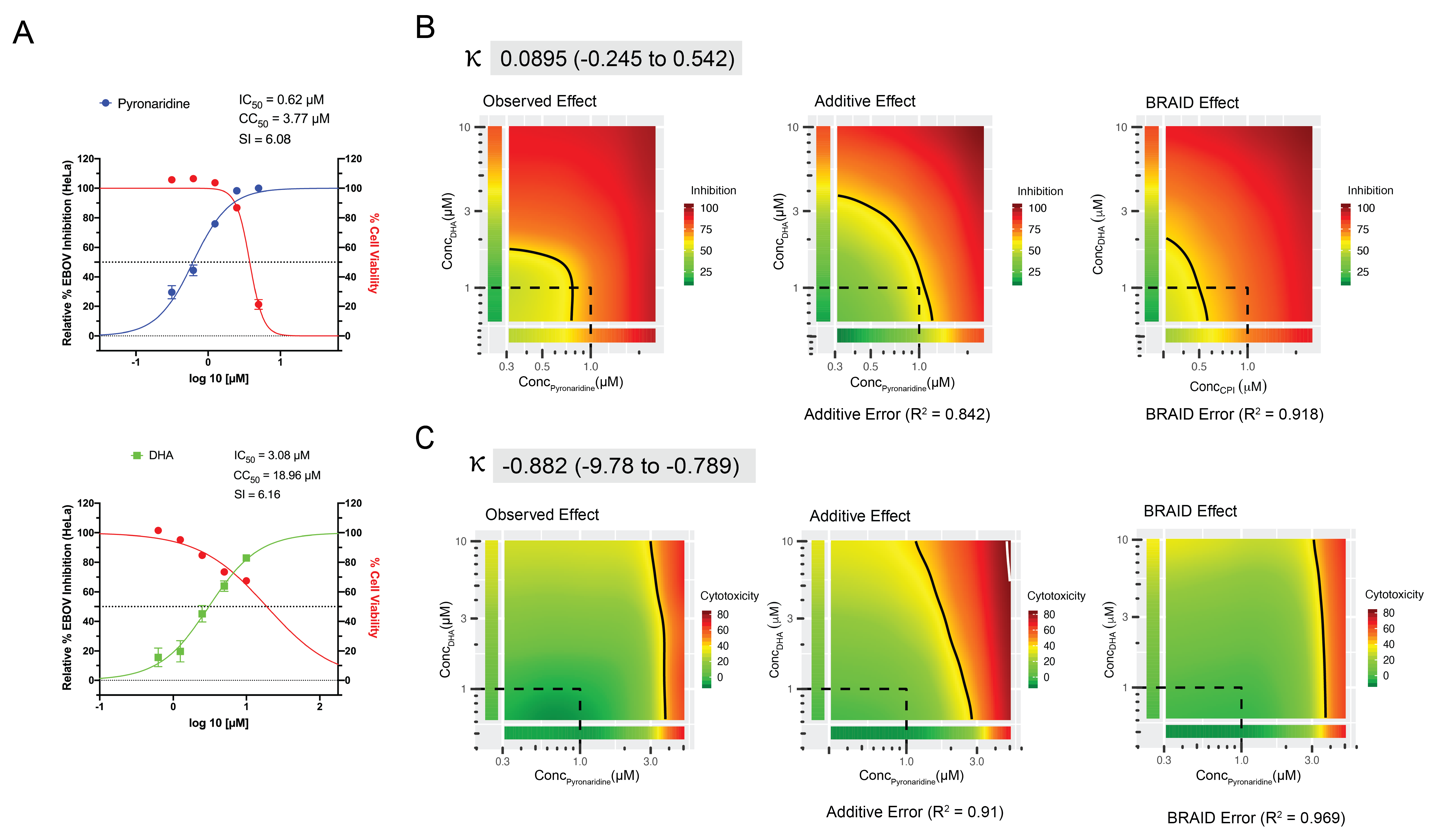

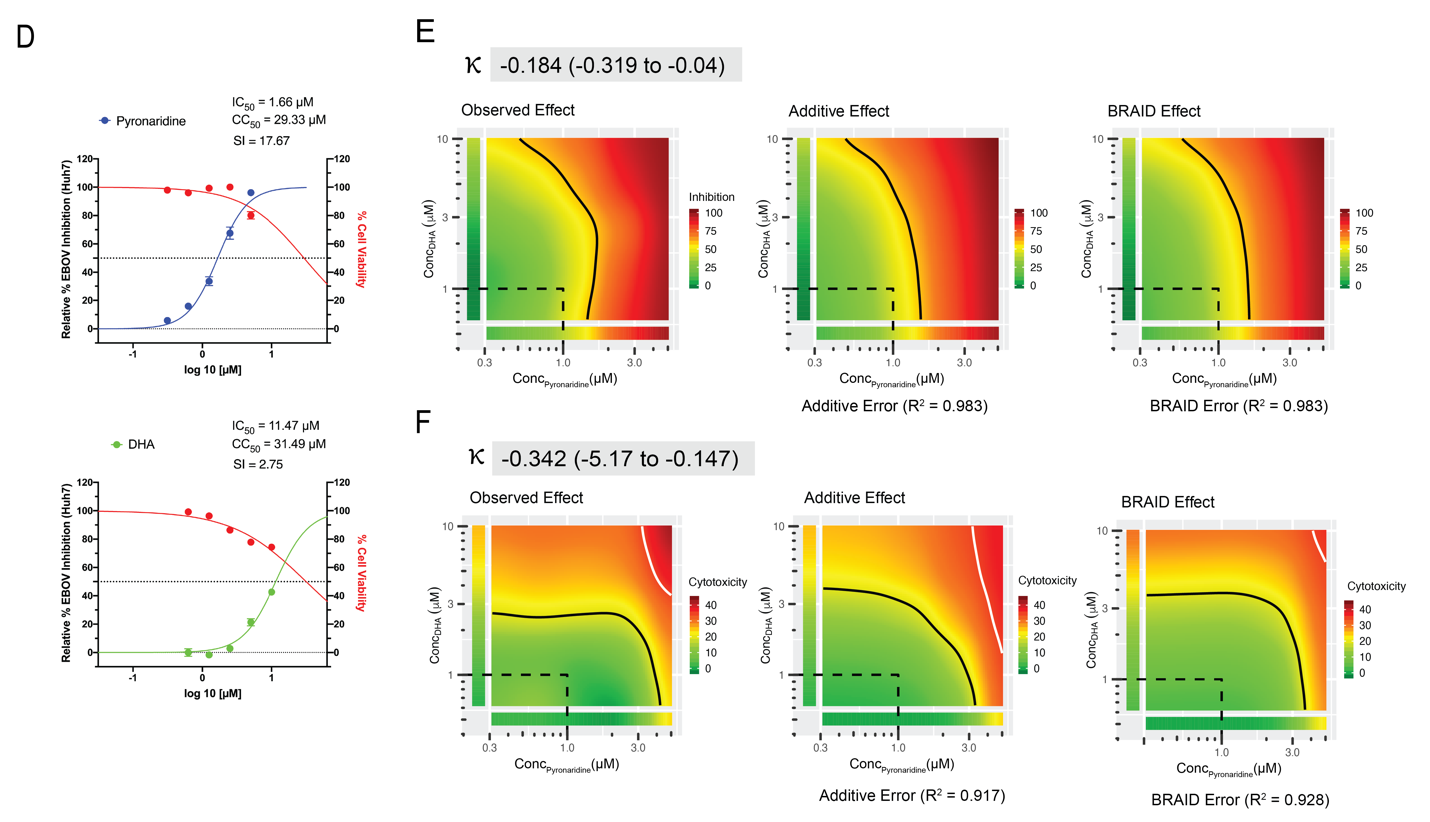

Figure S5. Combination data for the pyronaridine and artesunate checkerboard assay in HeLa cells. (A-B) Inhibition (top) or cell toxicity (bottom) matrixes (left) and 2D synergy maps (right). Synergy maps highlighting synergistic and antagonistic dose regions in red and green colors, respectively. The summary synergy score can be interpreted as the average excess or reduced response due to simultaneous drug treatments. A synergy score near 0 gives limited confidence on synergy or antagonism and a score < -10 or >10 are expected to be antagonist or synergistic, respectively. These analyses suggested that inhibition is additive, while cellular toxicity is antagonist when these compounds are given simultaneously.

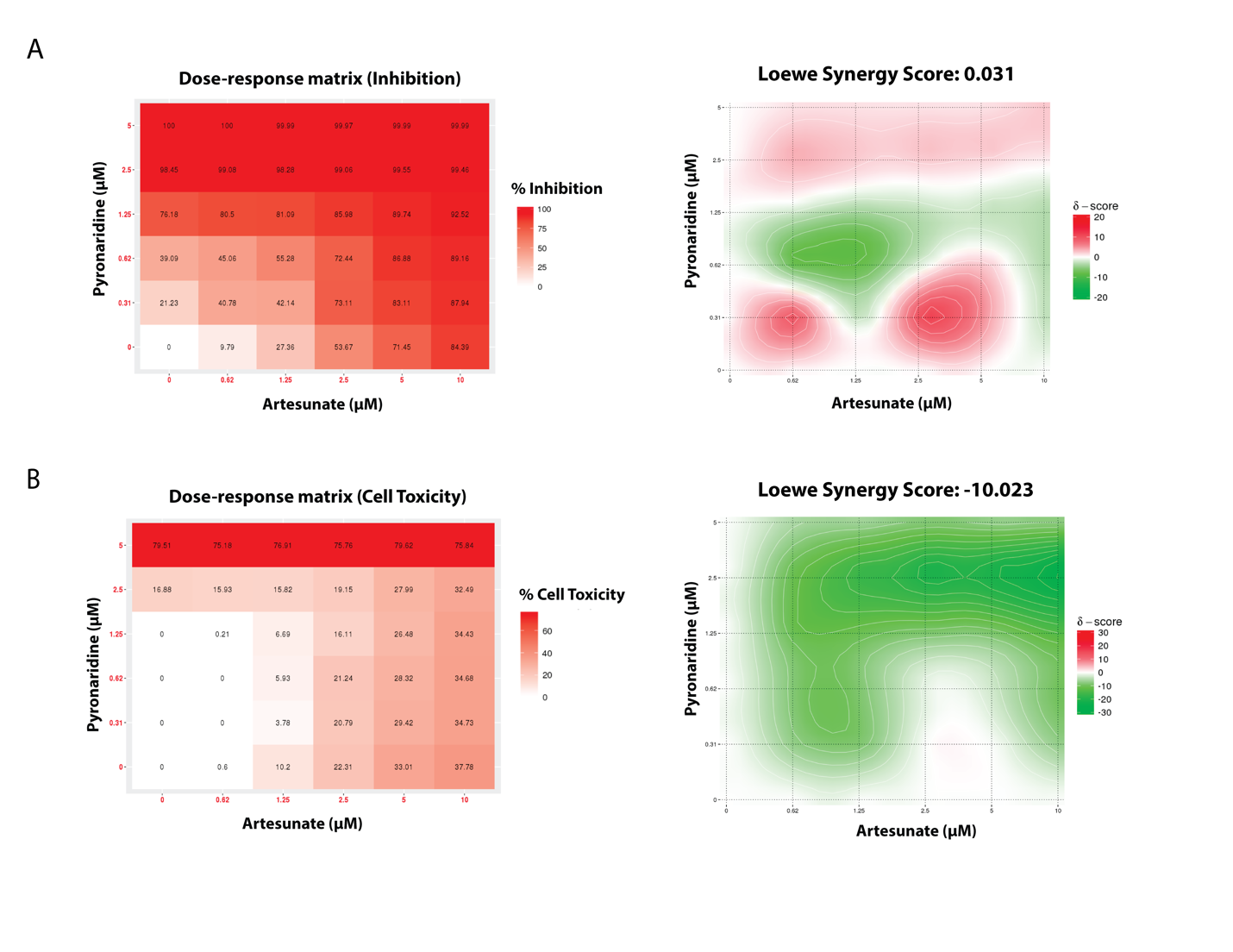

Figure S6. Combination data for the pyronaridine and DHA checkerboard assay in HeLa cells. (A-B) Inhibition (top) or cell toxicity (bottom) matrixes (left) and 2D synergy maps (right). Synergy maps highlighting synergistic and antagonistic dose regions in red and green colors, respectively. The summary synergy score can be interpreted as the average excess or reduced response due to simultaneous drug treatments. A synergy score near 0 gives limited confidence on synergy or antagonism and a score < -10 or >10 are expected to be antagonist or synergistic, respectively. These analyses suggested that inhibition is additive, while cellular toxicity is antagonist when these compounds are given simultaneously.

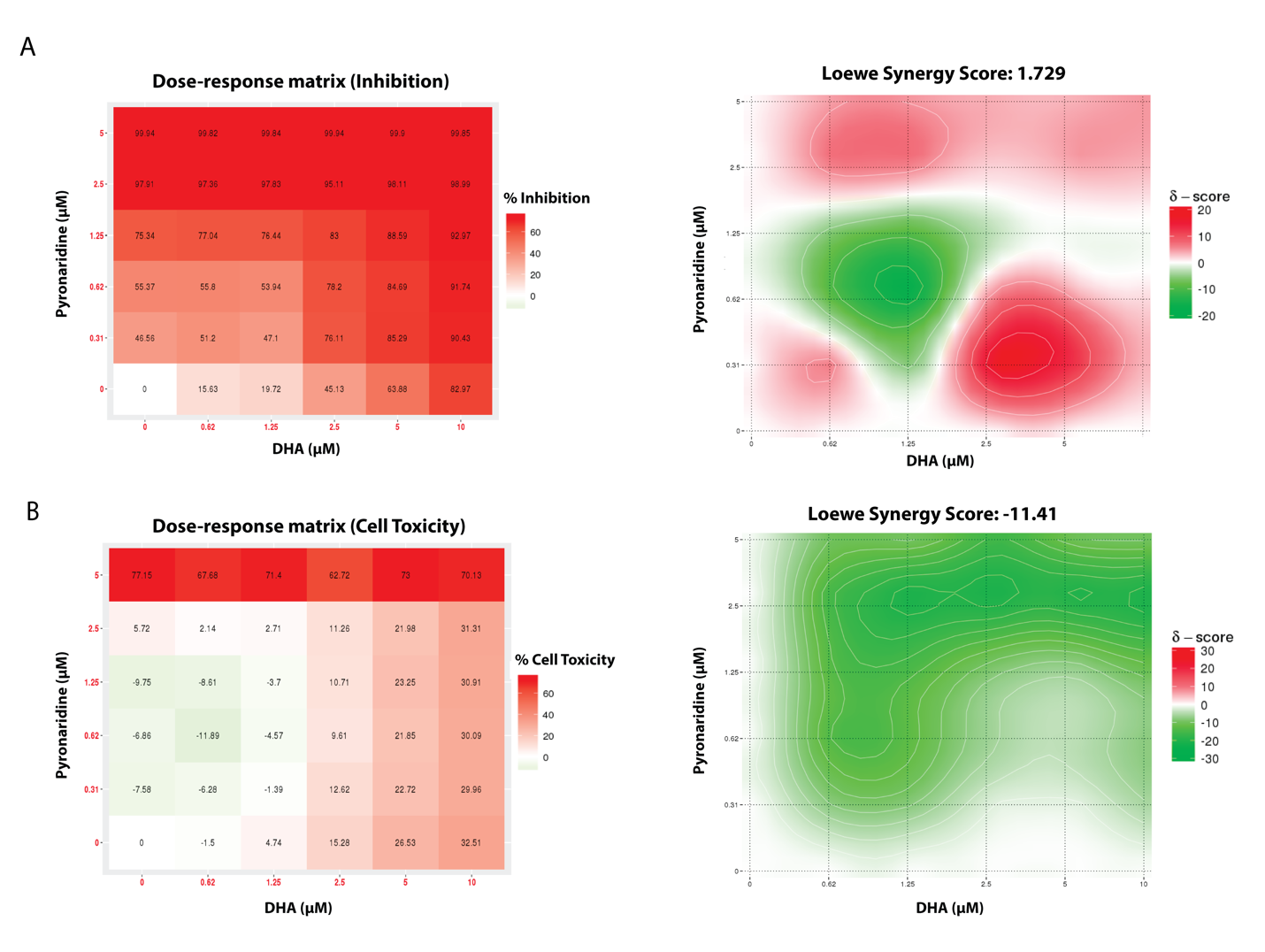

Figure S7. Combination data for the pyronaridine and DHA checkerboard assay in Huh7 cells. (A-B) Inhibition (top) or cell toxicity (bottom) matrixes (left) and 2D synergy maps (right). Synergy maps highlighting synergistic and antagonistic dose regions in red and green colors, respectively. The summary synergy score can be interpreted as the average excess or reduced response due to simultaneous drug treatments. A synergy score near 0 gives limited confidence on synergy or antagonism and a score < -10 or >10 are expected to be antagonist or synergistic, respectively. These analyses suggested that both inhibition and toxicity are either additive or mildly antagonist when these compounds are given simultaneously.

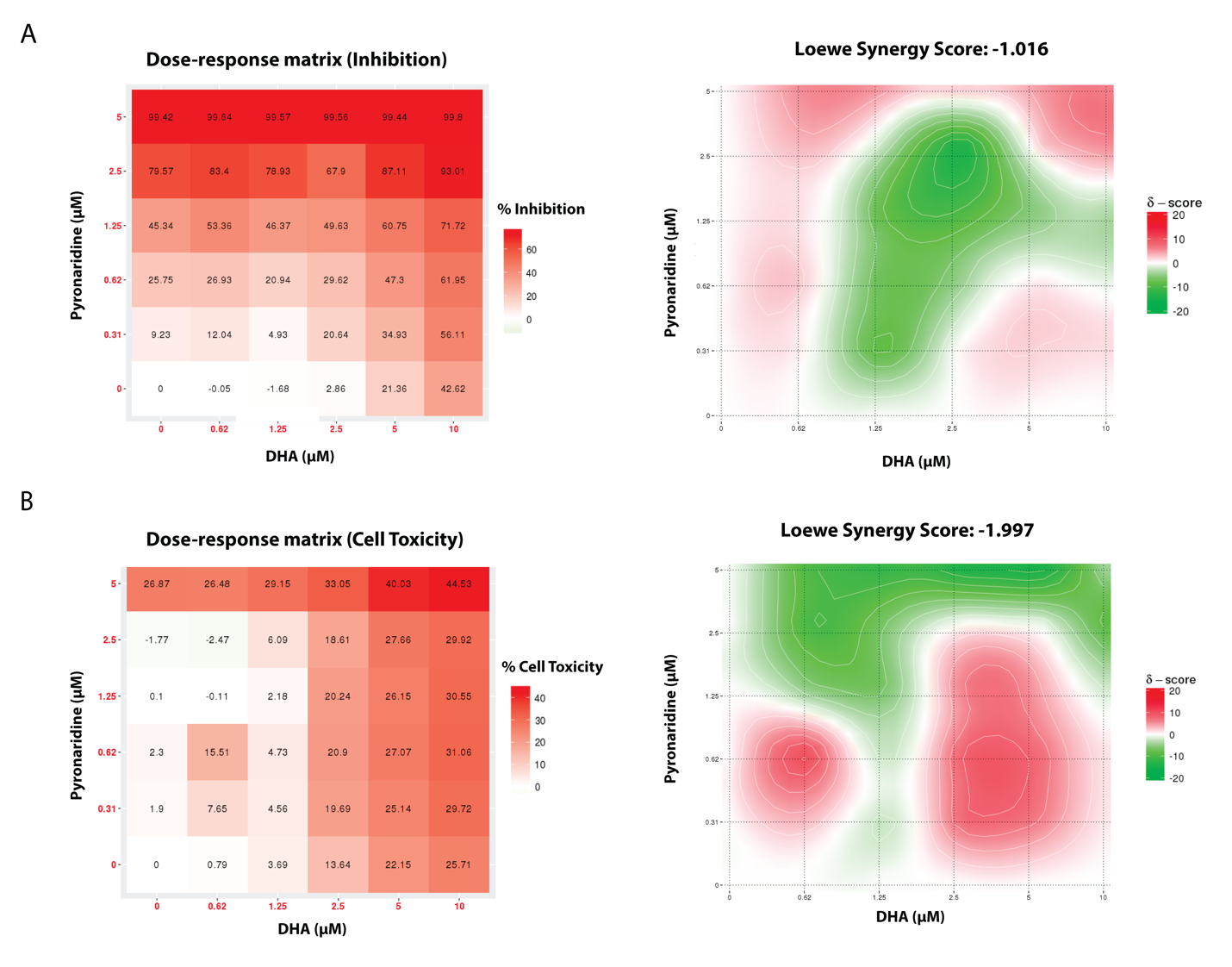
